# Label-free cleared tissue microscopy and machine learning for 3D histopathology of biomaterial implants

**DOI:** 10.1101/2022.11.16.516791

**Authors:** Tran B Ngo, Sabrina DeStefano, Jiamin Liu, Yijun Su, Hari Shroff, Harshad D Vishwasrao, Kaitlyn Sadtler

## Abstract

Tissue clearing of whole intact organs has enhanced imaging by enabling the exploration of tissue structure at a subcellular level in three-dimensional space. Although clearing and imaging of the whole organ have been used to study tissue biology, the microenvironment in which cells evolve to adapt to biomaterial implants or allografts in the body is poorly understood. Obtaining high-resolution information from complex cell-biomaterial interactions with volumetric landscapes represents a key challenge in the fields of biomaterials and regenerative medicine. To provide a new approach to examine how tissue responds to biomaterial implants, we apply cleared tissue light-sheet microscopy and three-dimensional reconstruction to utilize the wealth of autofluorescence information for visualizing and contrasting anatomical structures. This study demonstrates the adaptability of the clearing and imaging technique to provide sub-cellular resolution (0.6μm isotropic) 3D maps of various tissue types, using samples from fully intact peritoneal organs to volumetric muscle loss injury specimens, with and without biomaterials implants. We further apply computational-driven image classification to analyze the autofluorescence spectrum at multiple emission wavelengths to categorize tissue types in the tissue-biomaterial microenvironment.

## INTRODUCTION

A critical stage of developing biomaterials for implantation into a patient involves an evaluation of tissue structure at the implantation site to provide information such as scaffold integration, tissue regeneration, or any pathology from adverse responses. Frequently and most commonly, these evaluations occur in two dimensions through cross-sectional analysis using histopathology of excised tissue (*1*). Histopathology provides detailed resolution and sub-cellular structural information of tissues. The multitude of staining options –direct chemical stains and fluorescent probes– have generated a wealth of information in phenotyping the structure of samples in cross-section (*2*, *3*). Recent advances utilize machine learning to convert two-dimensional serial sections stained with hematoxylin and eosin (H&E) into 3D reconstructions and cleared tissue microscopy to visualize fluorescence in intact tissues (*4*–*6*).

Advances in light-sheet and cleared tissue microscopy have led to the development of threedimensional evaluations of different tissues, especially in neuroscience research (*7*–*11*). Lightsheet or selective plane illumination microscopes (SPIMs) are well suited for imaging large samples of cleared tissue due to their unmatched speed and confined illumination (*12*). At typical operating framerates of 10-100Hz, SPIMs are capable of imaging centimeter scale volumes within hours rather than days. In thick, densely labeled, or autofluorescent samples, restricting the illumination to only the image plane considerably decreases the out-of-focus fluorescence that would otherwise deteriorate spatial contrast. Specific implementations of SPIM, such as the DISPIM, can acquire multiple complementary views of the sample and computationally fuse them to improve resolution, reduce striping, and correct for attenuation at the cost of additional acquisition and computation time (*13*). While the DISPIM is capable of sub-micron isotropic resolution, not all applications demand it. The flexibility of the DISPIM enables us to select the acquisition/computation protocol that best suits the specific experimental need by balancing image quality against total pipeline time.

Different clearing protocols have been developed to improve in-depth light penetration through the whole-mount tissue. In cleared tissue imaging, autofluorescence has generally been regarded as a contaminating signal that must be minimized either chemically or spectrally. However, autofluorescence contains a wealth of structural contrast that can provide an anatomical context for targeted exogenous probes (*14*, *15*). Using a simple and inexpensive DISCO series clearing method and the cleared tissue dual-view inverted selective plane illumination microscope (ct-diSPIM) (*12*, *16*, *17*), we capture autofluorescence to visualize the different tissue structures in the intact mouse peritoneal cavity, including the liver, kidney, intestinal lining, along with a detailed 3-dimensional view of a decellularized extracellular matrix (ECM) biomaterial implant in a volumetric muscle loss injury.

Furthermore, autofluorescence in an empty channel, such as the green channel, allows us to visualize tissue-specific histological characteristics with immunolabeling. In this study, we have translated this principle into a three-dimensional reconstruction of muscle injury. The autofluorescence of muscle tissue at multiple emission wavelengths can be analyzed and interpreted through machine learning to differentiate from surrounding interstitium, blood vessels, and ECM biomaterials in the complex implantation microenvironment. We demonstrate the applications of label-free cleared tissue microscopy and machine learning based on iDISCO/3DISCO clearing methods for examining the three-dimensional histopathology of biomaterial implants.

## MATERIALS & METHODS

### Extracellular matrix decellularization

The small intestine from Yorkshire Pigs (Wagner Meats) was dissected to remove luminal and muscular layers to isolate the submucosa. The submucosa was rinsed thoroughly in distilled water with 1% antibiotic antimycotic and stored at −80 °C until processing. The tissue was then thawed and rinsed in sterile distilled water and then incubated in 4 % ethanol (Sigma) with 0.1 % peracetic acid (Sigma) diluted in sterile water for 30 minutes on a stir plate at room temperature. The resulting material was washed with sterile distilled water and phosphate-buffered saline (1xPBS) until the pH returned neutral. Samples were frozen overnight at −80 °C and then lyophilized for 3 days before being milled into a powder using a SPEX Cryogenic Milling device.

### Volumetric muscle loss

Female 6–8 week-old C57BL/6 mice (Jackson Labs) were anesthetized under isoflurane (2-4%) and subsequently given 1mg/kg buprenorphine SR (ZooPharm) for pain relief before surgery. The volumetric muscle loss (VML) was performed as previously published (*18*). Briefly, 24 hours before surgery, hair was removed with an electric razor and depilatory cream to clean an area from the hindfoot up to the xiphoid process. On the day of surgery, the area was sterilized with three successive scrubbings of betadine followed by 70% isopropanol. A 1 cm incision was made in the skin overlying the quadriceps muscle group, followed by the fascia, and the inguinal fat pad was pushed toward the hip joint to provide access to the quadriceps. A 3 mm portion of the quadriceps was removed using surgical scissors, and the resulting gap was filled with either a saline vehicle control or ECM powder hydrated in saline. The skin was closed using 3 – 4 wound clips before repeating the procedure on the contralateral leg. Mice were monitored under a heat lamp until fully recovered and ambulatory. All animal procedures were reviewed and approved by the NIH Clinical Center ACUC protocol number NIBIB 20-01.

### Tissue Collection

After 3-week post-implantation, studied animals were heavily anesthetized with 4% isoflurane and assured surgical-plane anesthesia by toe pinch. Then, the animals were subcutaneously injected with 100 mL heparinized PBS (10 U/mL) to prevent premature clotting. For perfusion, the initial incision was made below the xiphoid process, followed by two lateral cuts of the rib cage to the sternum providing access to the thoracic cavity. The right atrium was carefully lacerated, and heparinized saline was perfused into the left ventricle via a 25 G needle at a rate of 5 mL/min. Following perfusion, each tissue – quadriceps muscle groups or intact peritoneal cavity – was harvested with special care to remove skin and hair. Tissues were fixed in Bouin’s solution (Sigma) for 2 days at room temperature (peritoneal cavity) or in 4% paraformaldehyde (Electron Microscopy Sciences) overnight at 4 °C (muscle tissue).

### Peritoneal Specimen Preparation

The intact peritoneal cavity specimens were rinsed in distilled water thoroughly to remove residual Bouin’s fixative and coloration. Each was divided into 3 smaller sections of approximately 10 mm thickness using a single-edge razor blade for better maintenance and sectioning. These smaller sections were rinsed in distilled water to remove Bouin’s solution. The peritoneal blocks were then flushed with distilled water using a 3-cc syringe and an 18G blunt tip to remove excess waste in the gastrointestinal tract. Each tissue block was embedded in 3% warm agarose (Sigma) and cut into 1000-2000 μm thick sections using a vibratome (Leica VT1200). Alternatively, samples were kept intact in 10 mm sections for clearing and imaging.

### Tissue Clearing using IDISCO

Both peritoneal sections and quadricep femoris specimens were washed with distilled water three times before clearing. The tissues were dehydrated in successive incubations with water-to-methanol series. Then the clearing procedure was followed by a methanol-to-dichloromethane series (DCM, Sigma) (*16*). The final refractive index solvent was matched in two changes of dibenzyl ether (DBE, Sigma) before mounting on a clear glass slide with Norland Optical Adhesive 81 by exposing the specimen in adhesion to ultraviolet light for 30-60 seconds to harden. Each specimen was left to re-equilibrate in DBE overnight before imaging on the ct-DISPIM. If there were air bubble artifacts, samples were subject to vacuum pressure from a laboratory vacuum supply while submerged in DBE to degas the sample.

### Cleared tissue DISPIM

The Dual-view Inverted Selective Plane Illumination Microscope (DISPIM) (Applied Scientific Instrumentation; Eugene, OR) optimized explicitly for cleared tissue, and the associated data processing pipeline has been described in detail previously (*12*). Briefly, a pair of 0.7NA multiimmersion objectives (Special Optics; Denville, NJ) was used to acquire images of cleared tissue samples either in single or dual view mode. Image volumes were acquired as stage-scanned tiles at full frame (2048 x 2048 pixels, FOV 520 μm, 0.254 μm per pixel) with 1 μm perpendicular inter-plane distance and 15% overlap between adjacent tiles. The sample was excited by a digitally scanned 637nm OBIS laser (Coherent; Santa Clara, CA) light sheet, and fluorescence was filtered through a 676/37 bandpass filter (Semrock; West Henrietta, NY) before being recorded on a Hamamatsu Flash 4 v3 sCMOS camera (Hamamatsu Photonics; Shizuoka, Japan) with a 20ms exposure time. Image tiles were then stitched and deskewed on the NIH Biowulf supercomputer and either deconvolved in the case of single-view or coregistered and joint-deconvolved in the case of multiview data. Three-dimensional rendering was completed in Imaris (version 9.9, Oxford Instruments) after conversion of raw stiched and deconvolved data to Imaris file type.

### Histology

Samples were collected and fixed as per protocol for ct-DISPIM preparation. Subsequently, they were dehydrated in successive changes of a water-to-ethanol series followed by clearing with xylenes and embedding in paraffin wax. The resulting blocks were sectioned on a rotary microtome (Leica) at 5 μm thickness. Samples were mounted on charged slides and dried overnight in a 55°C oven. Slides were stained with hematoxylin and eosin (H&E, Sigma) per the manufacturer’s instructions. Slides were imaged on an EVOS microscope. For the whole peritoneal section, which exceeded the microscope’s field of view, views were stitched together using the stitching plugin in ImageJ (version 1.41) (*19*).

### Data analysis and statistics

Images shown are representative of *n* = 5 – 6 samples. Due to file size exceeding the size limit of uploads, two videos showing the full 3D reconstruction of one representative peritoneal cavity with visceral organs (**Supplemental Video 1**) and quadriceps muscle with scaffold implant (**Supplemental Video 2**). Full stitching, deconvolution, and 3D reconstruction was only completed on these samples, with *n* = 5 peritoneal samples and *n* = 6 muscle samples observed via generation of one Z-stack. Machine learning (image segmentation) was completed in MatLab R2022b using the imsegkmeans K-means clustering-based image segmentation algorithm. Twenty-four (24) Gabor filters were implemented to cluster tissue types based on various factors. See Results for more information.

## RESULTS

To render and visualize a large section of biomaterial implants in the native tissue, we modified a cost-effective and commonly used IDISCO clearing technique for light sheet imaging. We tested the application of the IDISCO clearing protocol to examine ECM implants in two different anatomical regions: in the peritoneal cavity of a fully intact murine body trunk and a muscular injury of a murine quadriceps femoris muscle. As the IDISCO protocol did not include a decalcification step, we used Bouin’s solution as a fixative and decalcification agent for the peritoneal specimens with the vertebrae columns to help with coarse sectioning of tissue into a manageable size. We perfused and flushed the tissue to remove blood and waste before clearing, then completed dehydration steps with a graded series of methanol, followed by a methanol-to-dichloromethane series (DCM) (**Figure 1**). The final refractive index solvent was matched in two changes of dibenzyl ether (DBE) before imaging on the ct-DISPIM while submerged in DBE. Additionally, as the large specimen requires a large volume of solvent for each step (5-10 mL per sample), the IDISCO protocol allows us to achieve clearing with costeffective reagents for both specimens. Without decalcification, the muscle tissue in the whole thigh specimen achieved complete transparency. By contrast, the femur bone in the same specimen did not reach transparency (**Figure 2**). With decalcification, a fully intact cross-section of the body trunk, including various visceral organs in the peritoneal cavity, reached transparency with the main artifact due to waste remnants in the gastrointestinal tract (**Figure 2**). Further clearing and removing any air-derived artifacts (bubbles) could be improved using a vacuum chamber after the sample was cleared in DBE to remove any remaining DCM and degas the sample.

**Figure 1:**
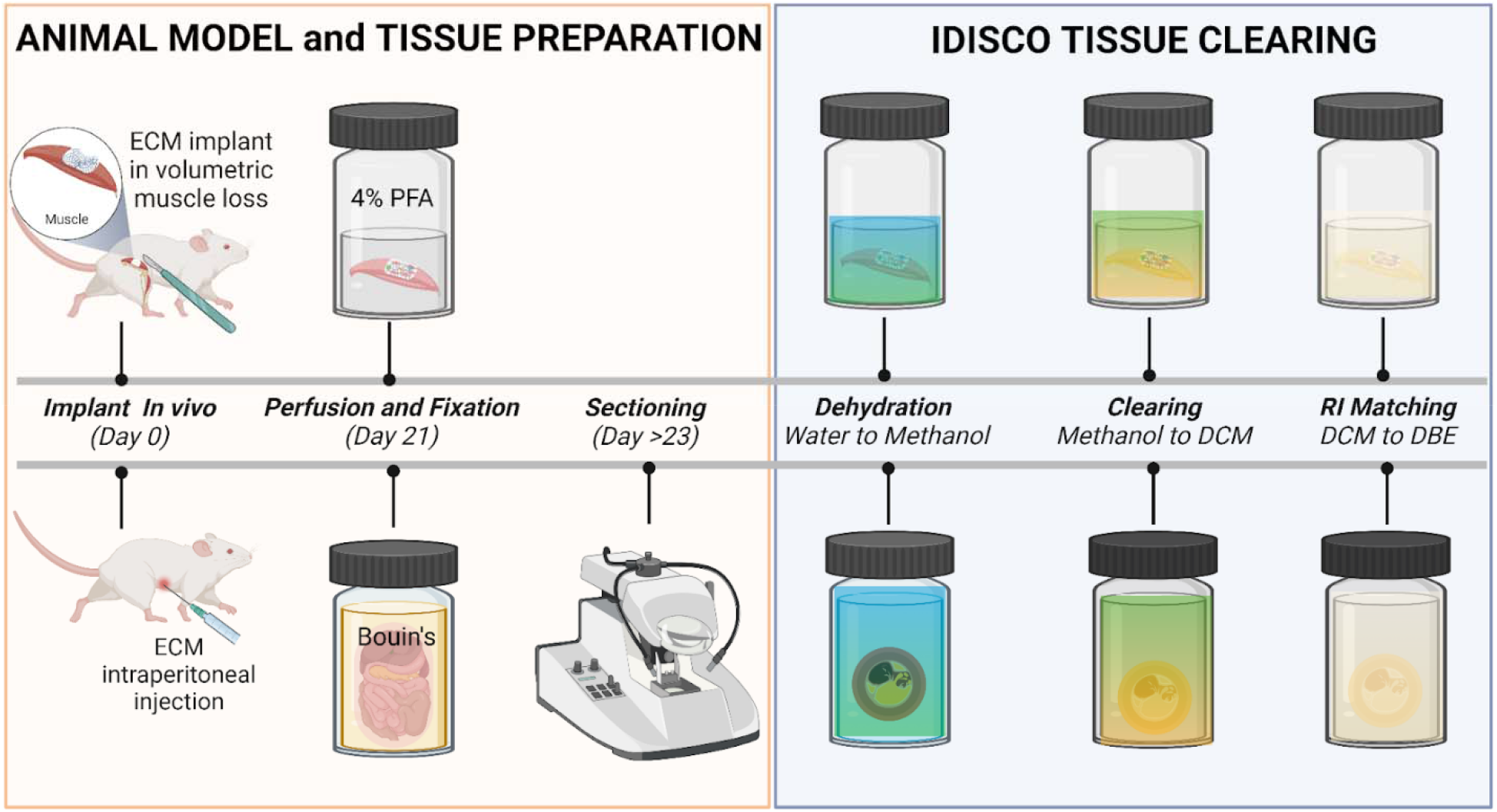
Animal model, tissue preparation, and IDISCO clearing protocol workflow. ECM biomaterials were implanted *in vivo* using two types of murine models: the volumetric muscle loss injury model and the intraperitoneal implant model. Specimens were collected and fixed on Day 21. The intraperitoneal specimens were cut into 1000-2000 μm sections using the vibratome or kept in whole 10 mm sections. The IDISCO clearing process for both tissue types starts with a graded water-to-methanol dehydration series followed by a 20% increasing graded methanol-to-dichloromethane (DCM) series before RI matching to dibenzyl ether (DBE) to achieve transparency. The specimen was incubated in each step of those graded series for at least 6 hours, which takes approximately 4 days for the entire clearing process. Figure created in BioRender.

**Figure 2:**
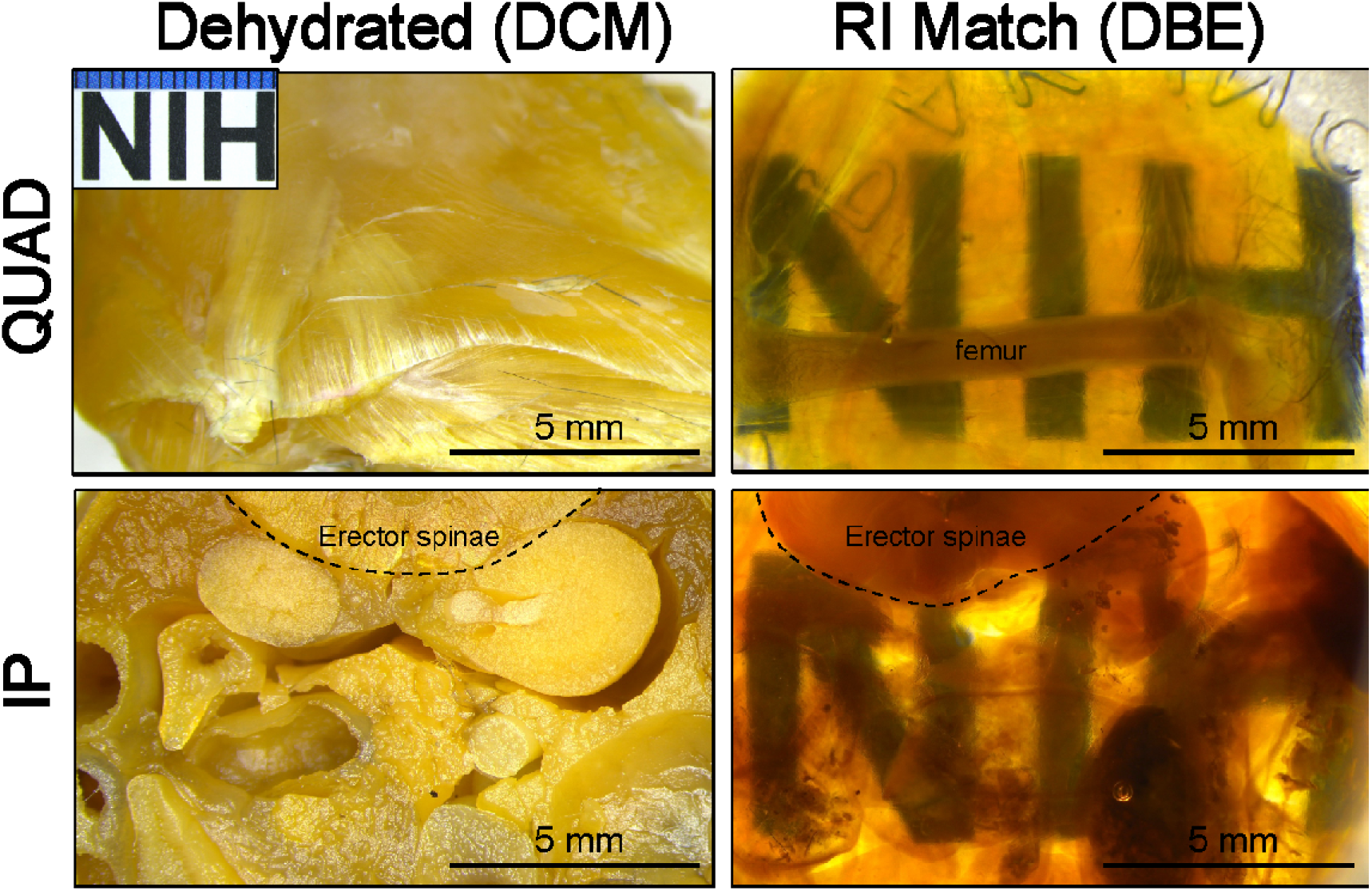
Stages of tissue clearing and refractive index matching in complex tissue structures. Tissues dehydrated in methanol followed by DCM are optically opaque. After refractive index (RI) matching in DBE, tissues are optically cleared. Quad = quadriceps muscle group, IP = peritoneal body cavity with visceral organs.

### Light-sheet imaging of large-volume tissue

We acquired image datasets of cleared tissue in DBE (RI: 1.562). At this refractive index (0.7NA, 676nm emission wavelength), the ct-DISPIM has a resolution of 0.6 μm laterally and 4.3 μm axially. This can be improved to 0.4 μm lateral and 3 μm axial with single-view deconvolution and 0.4 μm isotropic resolution with dual-view deconvolution. In our current work with the peritoneal specimen decalcified with Bouin’s solution, we acquired image volumes up to 2 mm in depth. Image resolution degraded as a function of depth due to imperfect clearing; however, a satisfactory (cellular) resolution could be achieved to a depth of 1 - 3 mm, depending on the sample.

While the ct-DISPIM is capable of 100Hz imaging, we found that operating at a slower 20ms framerate, with a total slice time of 45ms to allow for additional hardware/software operations, ensured better system stability for substantial data acquisitions (>1TB). At this slower framerate, acquiring a 1 mm long image stack (tile) with a 1 μm perpendicular inter-frame distance (707 slices) requires 32 seconds. Samples imaged in this work typically required 100-300 tiles to cover, each tile consisting of 1000-15,000 slices.

After deconvolution and stitching, a high-resolution Z-stack was generated that could be interrogated at a variety of magnifications (**Figure 3**). In the peritoneal cavity, the entire body cavity with visceral organs could be projected and bore a striking similarity to standard hematoxylin and eosin stained 5 μm section (**Figure 3a,d**). Spatial resolution was sufficient to allow us to visualize macroscopic (**Figure 3a,d**), microscopic multi-cellular (**Figure 3b,e**), and single cellular structures such as goblet cells in the intestine (**Figure 3c,f**). In a muscle injury model, the collagen-based ECM scaffold cleared completely. The tissue could be visualized in its entirety with minimal loss of information in comparison to an H&E stain (**Figure 3g,i**). As with the peritoneal cavity, high-resolution images could be obtained to interrogate the structure and cellular infiltration at the intersection of tissues, specifically the biomaterial-tissue interface, which is of interest to biomedical material research (**Figure 3h,i**).

**Figure 3:**
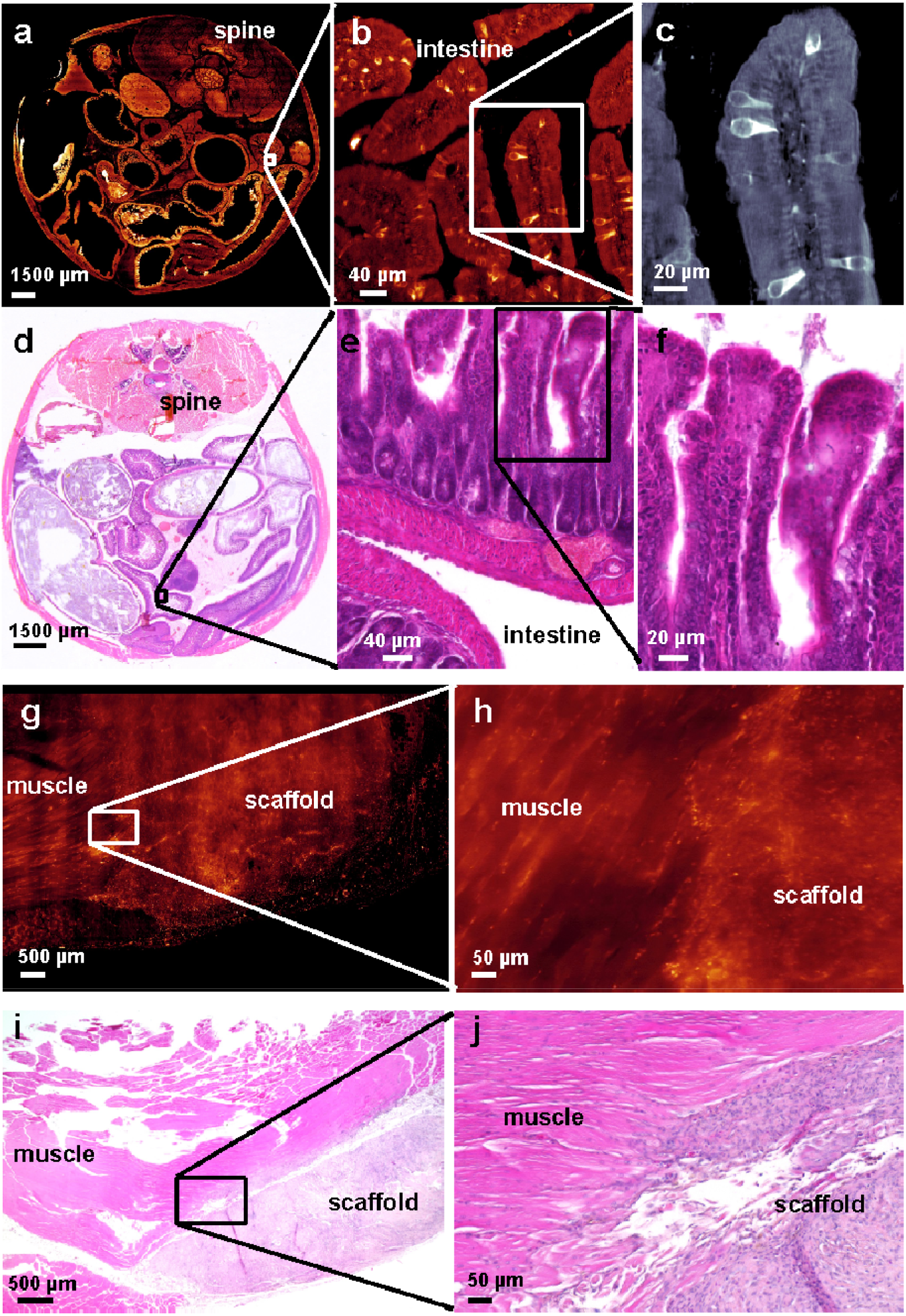
Range of resolution and tissue structures identified via ct-DISPIM using autofluorescence. **(a-c)** Images of the entire murine peritoneal section were acquired using autofluorescence (AF) excited at 637nm. (**d-f)** H&E stained 5 μm sections of a similar region in the mouse peritoneal cavity, focusing on the intestinal wall with similar magnifications to match a-c, respectively. (**g-h)** Higher magnification views of the intersection of the ECM scaffold in a volumetric muscle loss injury. (**i-j)** Representative H&E images of a similar tissue-biomaterial interface where the native muscle tissue integrates with the ECM scaffold at similar magnifications.

### Deep imaging of large intact multi-visceral organs in the peritoneal cavity and multi-tissue differentiation in a muscle injury model with high resolution

IDISCO clearing process preserves intact multi-visceral organs in the abdominal cavity (**Figure 4a-h**). As a result, we can visualize the prominent anatomical features of different organs in the abdominal cavity without immunolabeling (**Figure 4**). In the kidney, we could identify spherical renal corpuscles and tubules. In addition, we could identify several detailed renal features, including the Bowman’s capsule, proximal tubules, and collecting ducts. Compared to standard hematoxylin and eosin staining, these structures appear equivalent, if not more easily discernable with the dynamic range of autofluorescence emission(**Figure 4d,e**). We can also closely examine the other organs in the abdominal cavity, such as the lining of the intestinal wall (**Figure 4f,g**). In areas of dense hematoxylin staining, this area appears lighter (stronger autofluorescence signal) in the Z-stack from the ct-DISPIM. Furthermore, with the IDISCO clearing technique combined with the acid-based Bouin’s fixative solution, we also achieved transparency of the vertebral column (**Figure 4h,i**). When actively explored in 3 dimensions, these data show a wealth of information lost with standard 2D histopathology (**Supplementary Video 1**).

**Figure 4:**
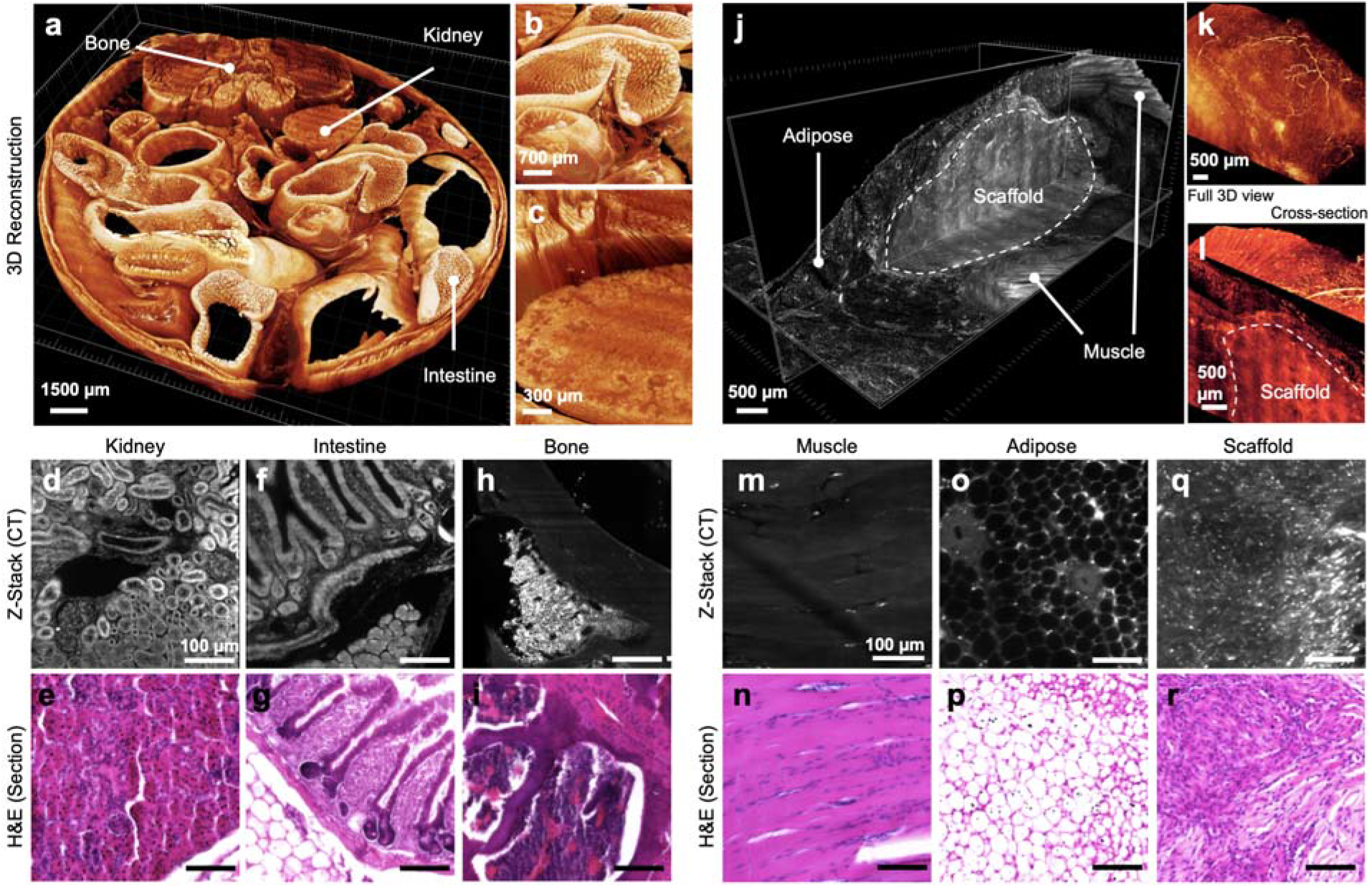
Three-dimensional reconstruction of multiple tissue types utilizing label-free ct-DISPIM. 3D projections of the peritoneal cavity and muscle injury with comparisons to standard H&E. **(a-c)** reconstruction of visceral organs in a fully intact abdominal cavity at different magnifications. (**a)** entire intact peritoneal cavity. (**b)** intestinal villi **c** kidney and erector spinae muscles. **(d-i)** tissue structures in comparison to corresponding 5 μm H&E-stained sections. (**d,e)** kidney (**f,g)** intestine (**h,i)** vertebra including bone and marrow. **(j-l)** muscle injury with biomaterial. **(j)** full 3D reconstruction of biomaterial implant **(k)** vascularization in overlaying fat pad. (**l**) muscle-implant interface. **(m-r)** tissue structures in comparison to corresponding 5 μm H&E-stained sections. (**m-n**) skeletal muscle, (**o-p**) adipose/fat pad (**q-r**) ECM scaffold.

In addition to the differentiation of very diverse solid organs within the abdominal cavity, we can also precisely evaluate the three-dimensional structures of biomaterial-tissue interactions within a volumetric muscle loss injury treated with an ECM scaffold (Figure 4 j-r). Specifically, we observed an adipose layer migrate over the scaffold material (**Figure 4j**). Furthermore, a strong vascularization of this adipose is seen overlying the scaffold (**Figure 4k**). The directionality of muscle fibers surrounding the scaffold can be seen through a combined cross-section and 3D projection of the tissue that could be leveraged in the future for quantitative comparison of outcomes in force generation with a linearity of fiber regeneration (**Figure 4l**). As with the peritoneal cavity, these different tissues can be compared to standard hematoxylin and eosin staining, wherein all of the same structures in muscle (**Figure 4m,n**), adipose (**Figure 4o,p**), and the ECM scaffold (Figure 4 q,r) can be differentiated and interrogated on a 3-dimensional basis (**Supplementary Video 2**).

### Utilizing autofluorescence to computationally reconstruct biomaterial-tissue interactions

Image segmentation is the classification of an image into different groups. Many kinds of research have been done in image segmentation using clustering. One of the most popular methods is K-Means clustering algorithm (*20*). K-Means clustering algorithm is an unsupervised algorithm that clusters or partitions the given image into K-clusters or parts based on the K-centroids of the feature set. The algorithm aims to find K groups based on similarity in the data (image pixels). The objective of K-Means clustering is to minimize the sum of squared distances between all data (image pixels) and the cluster center. The method steps include (1) choosing the number of clusters K; (2) randomly selecting K points as the centroids; (3) assigning each data point (image pixel) to the closest centroid; (4) computing and placing the new centroid of each cluster; (5) reassigning each data point (image pixel) to the new closest centroid. If any reassignment occurs, go to step (4); otherwise, the clustering is ready.

In this work, we use the K-Means clustering algorithm to cluster 4 different regions (Muscle, ECM Implants, Interstitial Space, and Blood Vessel) in the image. Our feature set included image intensity, texture information, and spatial information. A set of Gabor filters (24 Gabor filters, covering 6 wavelengths and 4 orientations) were used to compute the texture information (*21*). The coordinates of image pixels were used to represent the spatial information. The spatial location allows the K-means clustering algorithm to prefer groupings that are close together spatially. When this information is applied to multiple slices within a Z-stack, we can create a three-dimensional projection of label-free tissue classification (**Figure 6**).

**Figure 5:**
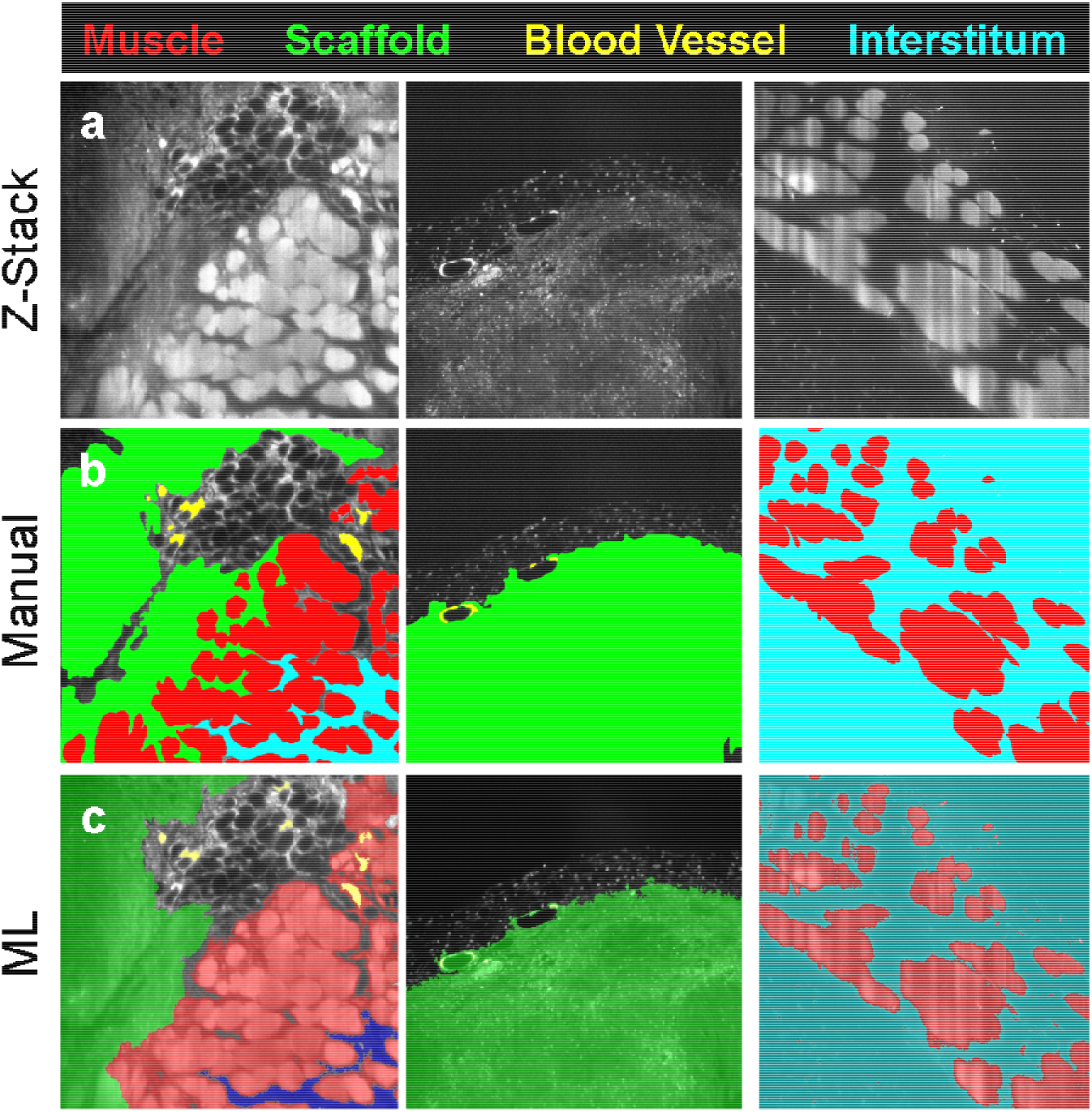
Machine learning development compared to manual tissue classification. (**a)** Example Z-stack images from cleared tissue imaging. (b) Manual classification of tissue types. (c) Machine learning (ML) outcome of tissue classification based on autofluorescence spectra and K-Means clustering. Red = muscle, Green = scaffold, Yellow = blood vessel, Blue = interstitium.

**Figure 6:**
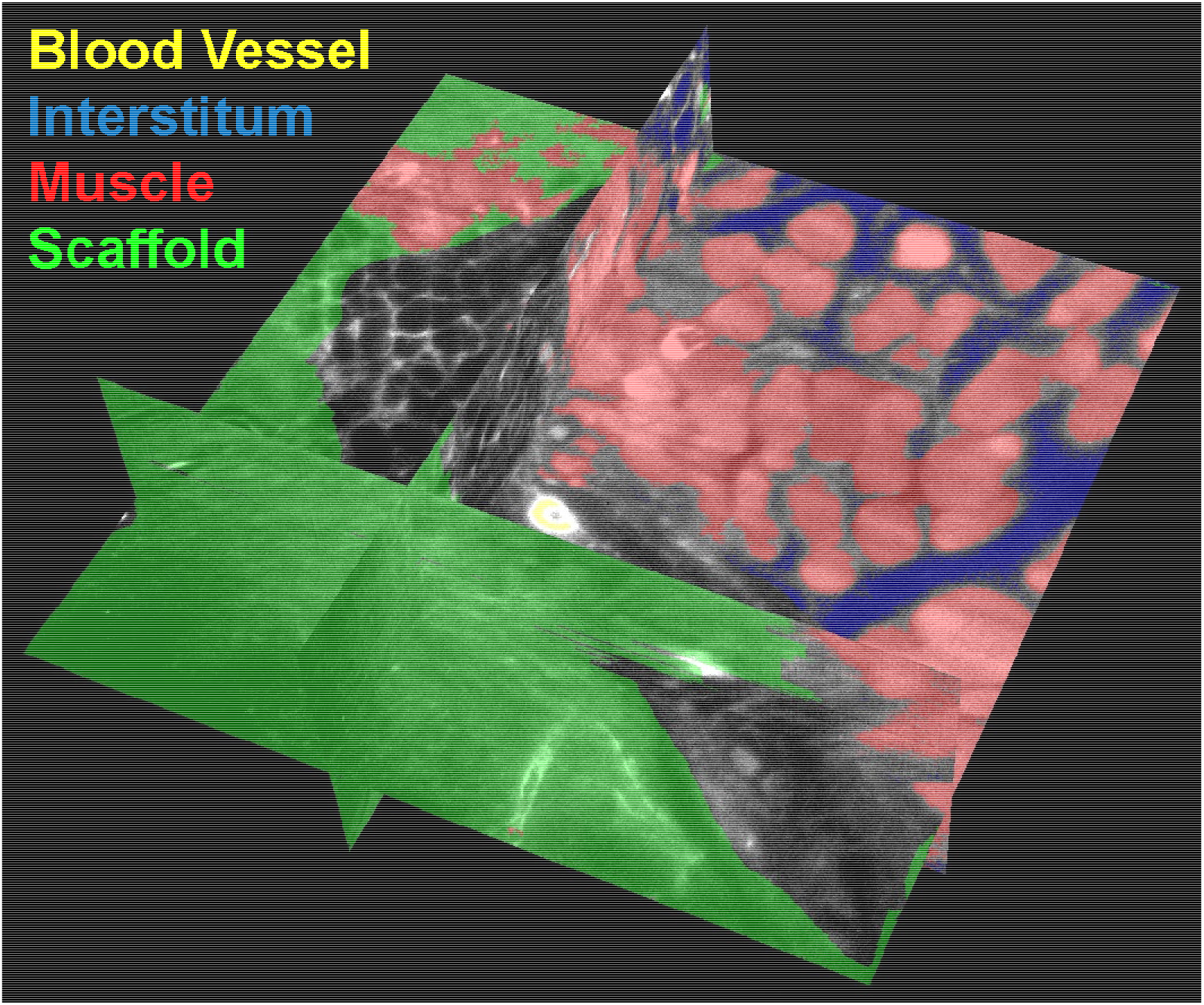
3D reconstruction of injury interface with machine learning classification of tissue classes. Projection showing the X-Y-Z axes of machine-learning-derived tissue classification overlaid on raw data from the scaffold-muscle interface. Yellow = blood vessel, blue = interstitium, red = muscle, green = scaffold.

## DISCUSSION

Tissue clearing, a technique of tissue preservation in which a solvent medium replaces water and lipids in biological tissues with a matching refractive index (RI) to achieve transparency, has been widely used to study cell interactions in their natural tissue settings. Combined with lightsheet microscopy, cleared tissues can be visualized macroscopically and/or analyzed at a cellular level while preserving the 3-D tissue complexity. Many clearing protocols were developed using various options for a medium, from organic solvent to a hydrogel system, to achieve optimal tissue transparency for deep imaging. Each clearing technique represents a practical approach to addressing a different research question. For example, iDISCO or Ce3D is useful for immunolabeling applications; CLARITY or CUBIC is desirable for preserving endogenous fluorescence of reporters such as GFP or tdTomato; PEGASOS aims to achieve transparency for nearly all tissue types, including long bone and teeth (*16*, *17*, *22*–*24*). As all clearing techniques aim to achieve high preservation of fluorescent signal, autofluorescence is often reported as noise interfering with target immunolabeling or endogenous fluorescence. However, the autofluorescence signature obtained at several excitation wavelengths can help identify different structures, such as to study skin pathology (*14*). Leveraging the wealth of information on autofluorescence, this study presents a new approach to utilizing light-sheet microscopy and machine learning to differentiate autofluorescent signals for investigating biomaterial-tissue interfaces.

The imaging depth is determined by tissue transparency and the working distance of microscope objectives. However, matching RI to achieve complete transparency of a large specimen containing multi-visceral organs is challenging due to various tissue types, hydroxyapatite crystals in bone, and/or debris. In this study, the desired transparency of the large specimen was achieved with iDISCO, a simple, rapid, and inexpensive method. iDISCO permits whole-mount immunolabeling with volume imaging of large cleared samples ranging from whole quadriceps muscle group to a fully intact section of the peritoneal cavity of an adult mouse. Our study utilized Bouin’s solution with the iDISCO protocol for the peritoneal cavity containing the vertebral column as a fixative and decalcification agent. Other clearing protocol, such as PEGASOS, has been attributed to include the decalcification step (*24*, *25*). Tissues fixed with Bouin’s solution have remnant yellow coloration that enhances the autofluorescence signal. However, Rindone *et al*. have demonstrated adequate bone tissue clearing transparency without decalcification (*8*).

iDISCO clearing method has its limitations when imaging the biomaterial-tissue interface. We found that the clearing organic solvent (DCM) in the iDISCO protocol did not preserve the fluorescent immunolabels of interest, which was not included in the validated antibodies list (*16*). However, the protocol is an excellent choice for preserving autofluorescence. In addition, DCM treatment can lead to differential shrinkage among soft and hard tissues, which was shown as interstitial space between muscle fibers. Lastly, applying iDISCO to clear non-natural derived biomaterials such as synthetic polymers or metals remains challenging. On the other hand, Yating *et al*. have applied PEGASOS to clear tissues with titanium and stainless steel implants (*25*). Therefore, we recommend selecting a clearing protocol specific to the material of interest and research questions and considering the volume size and cost of each clearing protocol.

The cleared tissues were stored and imaged in DBE with an RI of 1.562 with the ct-DISPIM. However, while the ct-DISPIM is capable of 100Hz imaging, we found that operating at a slower 20ms framerate, with a total slice time of 45ms to allow for additional hardware/software operations. At this slower framerate, acquiring a 1mm long image stack (tile) with a 1 μm perpendicular inter-frame distance (707 slices) requires 32 seconds. Due to the fact that imaging each sample typically requires 100-300 tiles to cover, with each tile consisting of 1000-15,000 slices, this process can accumulate substantial data acquisitions (>1TB) over >24 hours of acquisition time. Even though the imaging process requires a long acquisition time, our platform results in impressive image quality at this refractive index (0.7NA, 676nm emission wavelength). The ct-DISPIM has a resolution of 0.6μm laterally and 4.3μm axially, which can be improved to 0.4 μm and 3 μm with single-view deconvolution, respectively, and 0.4 μm isotropic resolution with dual-view deconvolution. With a satisfactory (cellular) resolution that could be achieved to a depth of 1-1.5mm, this clearing and imaging technique is suitable for capturing the 3D tissue-biomaterial interfaces.

In this study, we applied the simple and inexpensive IDISCO tissue-clearing method for studying biomaterial implant interactions in various tissue contexts. First, we demonstrated the application of IDISCO in optically clearing and imaging multi-visceral organs in the whole intact murine peritoneal cavity section, with and without biomaterial implants. Furthermore, we also applied this approach to study tissue-biomaterial interaction in volumetric muscle loss injury. Leveraging the wealth of autofluorescence and machine learning, we established an image segmentation pipeline to differentiate tissue and biomaterials compartments at the implantation sites. Overall, we showcased the broad application of IDISCO tissue clearing–based deep imaging as a valuable tool for studying tissue interactions with biomaterials in tissue engineering and regeneration.

## CONCLUSION

This study shows the applications of light-sheet microscopy to 3D-image optically cleared tissues by IDISCO to investigate biomaterial-tissue integration at the implantation site. The natural autofluorescence of cleared tissues enhances the volumetric reconstruction and visualization of different organs in the murine GI tract at both the macroscopic and cellular levels with a high resolution of 0.6 μm (isotropic). Furthermore, this imaging technique of cleared tissue can be applied to study biomaterials in treating volumetric muscle loss injury. The autofluorescence signature of cleared muscle tissue was utilized for image segmentation to differentiate native muscle tissues and vasculature from the implanted ECM biomaterials. Overall, this work brings a new approach to examining the biomaterial-tissue integration to understand the interface morphology and analyze morphological changes associated with regeneration or fibrosis.

## ACNKOWLEDGEMENTS

The authors would like to thank Dr. Ted Usdin for his guidance on sample preparation. This work was funded by the intramural research program of the National Institutes of Health, National Institute of Biomedical Imaging and Bioengineering, National Institutes of Health (NIH). This work used the computational resources of the NIH HPC Biowulf cluster (http://hpc.nih.gov). This work was supported by the Howard Hughes Medical Institute (HS). This article is subject to HHMI’s Open Access to Publications policy. HHMI laboratory heads have previously granted a nonexclusive CC BY 4.0 license to the public and a sublicensable license to HHMI in their research articles. Pursuant to those licenses, the author-accepted manuscript of this article can be made freely available under a CC BY 4.0 license immediately on publication. <u>Disclaimer</u>: The contents of this publication are the sole responsibility of the authors and do not necessarily reflect the views, opinions, or policies of the NIH and the Department of Health and Human Services (HHS). Mention of trade names, commercial products, or organizations does not imply endorsement by the U.S. Government.

## CONFLICT OF INTEREST

The authors declare no conflict of interest.

## DATA AVAILABILITY

Due to large file sizes of images in the multi-terabyte range, raw data and image files are available upon request. Due to limitations in file size upload to bioRxiv, Supplementary Video 2 is available via request.

## Notes

### Competing Interest Statement

The authors have declared no competing interest.

